# AMP accumulation during mitochondrial stress induces transcription of cytosolic and mitochondrial protein synthesis components via NHR-180

**DOI:** 10.1101/2025.01.22.634344

**Authors:** Avijit Mallick, Theresa Ramalho, Yunguang Du, Rui Li, Levi Ali, Sookyung Kim, Pooja Rai, Andreas Bergmann, Lihua Julie Zhu, Cole M. Haynes

**Affiliations:** Department of Molecular, Cell, and Cancer Biology University of Massachusetts Chan Medical School, Worcester, MA 01605, USA

**Keywords:** NHR-180, ATFS-1, mitochondria, OXPHOS, mtDNA, protein synthesis

## Abstract

The mitochondrial unfolded protein response (UPR^mt^), which promotes mitochondrial proteostasis and biogenesis is regulated by the transcription factor ATFS-1. In *C. elegans*, impaired *atfs-1* expression perturbs mitochondrial function and slows development. Here, via an RNAi suppressor screen, we found that inhibition of the nuclear hormone receptor NHR-180 increases mitochondrial function and the rate of development in *atfs-1(null)* worms. NHR-180 is activated during mitochondrial dysfunction and induces transcription of genes required for protein synthesis on cytosolic and mitochondrial ribosomes. Importantly, the NHR-180 ligand binding domain interacts with AMP, which results in increased protein synthesis potentially to promote mitochondrial biogenesis and recovery from transient mitochondrial perturbations. However, NHR-180 exacerbates the mitochondrial dysfunction caused by mutations in genes required for oxidative phosphorylation. Consistent with these findings, we found that inhibition of the S6 kinase homolog RSKS-1 also increased mitochondrial function in *atfs-1(null)* worms. Importantly, treatment with an S6 kinase inhibitor also increased respiration and mtDNA content in mammalian ATF5-knockout cells and OXPHOS-deficient models, suggesting that translation regulation is a conserved, targetable approach for restoring mitochondrial function in disease.

## Introduction

Mitochondria are double membrane-bound organelles located in the cytosol. They are required for essential cellular processes, including oxidative phosphorylation, nucleotide and amino acid synthesis, as well as regulating programmed cell death and innate immune signaling (Anderson and Haynes, 2020; Higuchi-Sanabria et al., 2018; Shpilka and Haynes, 2018). Mitochondrial dysfunction occurs in numerous diseases, including Parkinson’s disease and Alzheimer’s disease (Irvine et al., 2008; Wang et al., 2020) as well as those diseases caused by mutations in genes encoding essential mitochondrial proteins (Moraes et al., 1989). The majority of mitochondrial proteins are encoded by nuclear genes, translated on cytosolic ribosomes, and subsequently imported into mitochondria. However, in *C. elegans,* 12 essential OXPHOS proteins are encoded by the mitochondrial genome (mtDNA) located within the mitochondrial matrix (13 in mammals). Most mitochondrial-localized proteins harbor an amino-terminal mitochondrial-targeting sequence (MTS) required for import into mitochondria via import channels within the mitochondrial outer membrane and the inner membrane.

In response to mitochondrial perturbations, the mitochondrial unfolded protein response (UPR^mt^) is activated, resulting in increased transcription of genes required for mitochondrial biogenesis and to maintain mitochondrial proteostasis. In *C. elegans,* the pathway is regulated by the bZIP protein ATFS-1, while ATF5 and ATF4 regulate the UPR^mt^ in mammals (Fiorese et al., 2016; Nargund et al., 2012, 2015; Shpilka and Haynes, 2018; Shpilka et al., 2021). When the mitochondrial population is functional, most of the ATFS-1 is efficiently imported into the mitochondrial matrix, where it is degraded by the protease LONP-1 (Anderson and Haynes, 2020; Nargund et al., 2012; Shpilka and Haynes, 2018). In contrast, upon mitochondrial perturbations including OXPHOS dysfunction or impaired mitochondrial proteostasis, a fraction of ATFS-1 fails to be imported into mitochondria and traffics to the nucleus to induce the UPR^mt^ (Pellegrino et al., 2014; Uma Naresh et al., 2022; Wu et al., 2018).

Here, we sought to identify transcription factor(s) that promote mitochondrial function. Nuclear hormone receptors (NHRs) in *C. elegans* are a large family of transcription factors that regulate gene expression in response to developmental, metabolic, and environmental cues. We have identified the previously unstudied transcription factor NHR-180, which promotes transcription of genes required for protein synthesis on mitochondrial ribosomes as well as on cytosolic ribosomes. NHR-180-dependent transcription is induced upon AMP binding the ligand binding domain within NHR-180, which occurs during OXPHOS perturbation. Interestingly, inhibition of NHR-180 expression alleviates mitochondrial dysfunction in both OXPHOS-deficient *C. elegans* and mammalian cells. Importantly, S6 kinase inhibition also alleviates mitochondria dysfunction. This work reveals a fundamental and conserved mechanism by which metazoans reprioritize translational resources to preserve mitochondrial function, offering new insights into the coordination of cellular metabolism, development, and survival under stress.

## Results

### Identification of transcription factors that impact mitochondrial function

The bZIP transcription factor ATFS-1 is required to activate the UPR^mt^ in *C. elegans,* which includes the induced transcription of genes required for mitochondrial function, such as the molecular chaperones *hsp-60* and *mthsp-70* (*hsp-6* in *C. elegans*) that reside within the mitochondrial matrix (Lin et al., 2016; Nargund et al., 2012, 2015; Pellegrino et al., 2014; Shpilka and Haynes, 2018). Worms lacking *atfs-1* have severe mitochondrial dysfunction, abnormal mitochondrial morphology, reduced mtDNA content, increased susceptibility to oxidative stress and infection, and a developmental delay (Pellegrino et al., 2014; Wu et al., 2018; Yang et al., 2022). However, despite the delay, *atfs-1(null)* worms develop into mature adults.

Here, we sought to identify transcription factors that impact mitochondrial function and the rate of development in the absence of ATFS-1. To identify transcription factors that are differentially expressed in *atfs-1(null)* worms relative to wildtype worms, we queried our previously published RNA-sequencing data (Shpilka et al., 2021). We identified 122 genes that encode transcription factors that are differentially expressed in *atfs-1(null)* worms relative to wildtype worms. Each of the 122 genes was knocked down via RNAi in *atfs-1(null)* worms, and the time it took each strain to reach adulthood was evaluated (Supplementary Table 1). Ten genes were identified that increased the development rate of *atfs-1(null)* worms, including *nhr-2, tbx-9, lsl-1, ceh*-*49,* or *nhr-180* (Fig. 1A-C). While, mRNA levels of most of the 122 transcription factors were increased in *atfs-1(null)* worms; *nhr-180* transcripts were reduced in *atfs-1(null)* worms relative to wildtype worms, suggesting that *atfs-1* positively regulates transcription of *nhr-180* (Shpilka et al., 2021).

**Figure 1:**
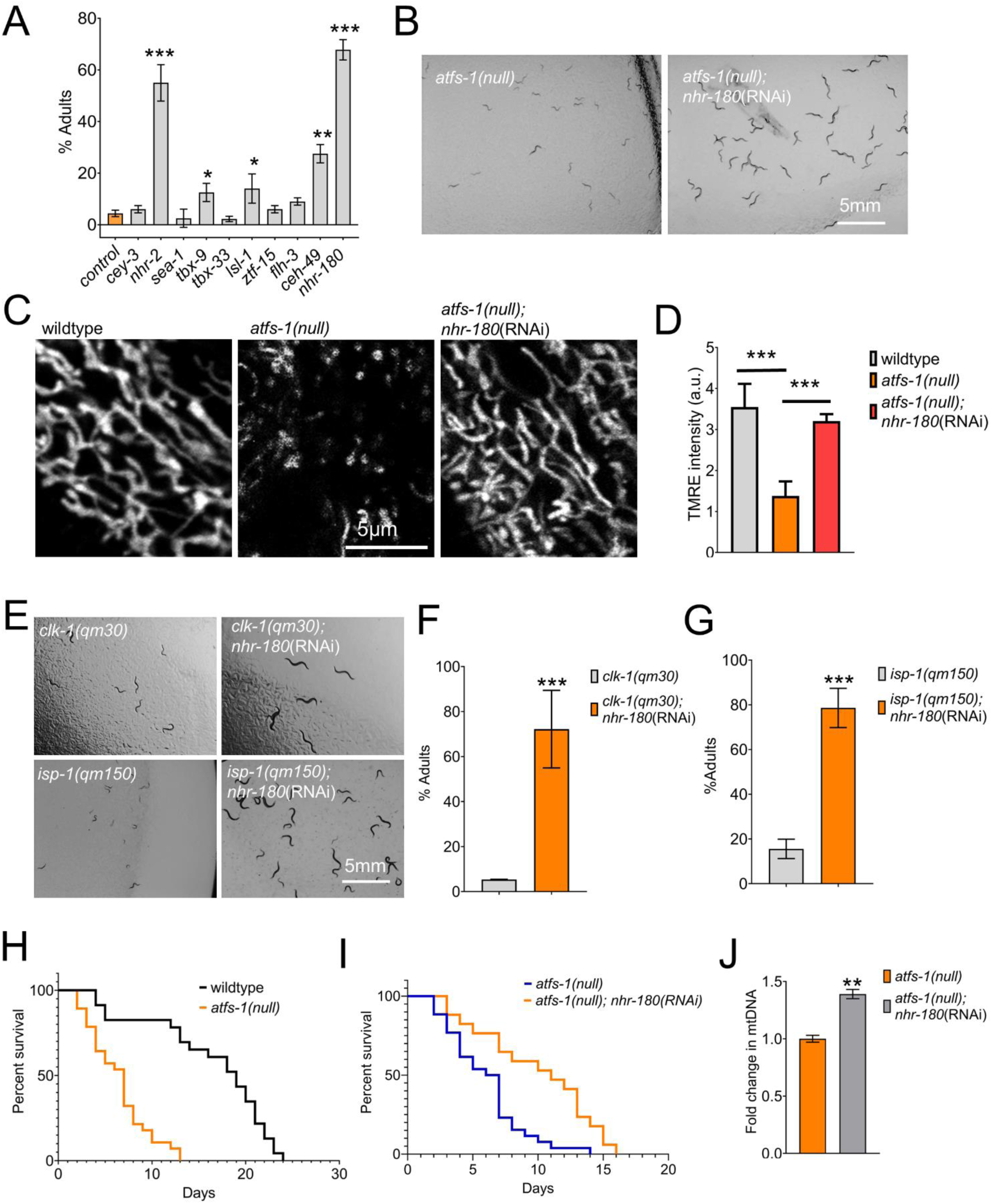
Inhibition of NHR-2 and NHR-180 suppresses slow growth in *atfs-1* null mutants. **A,B** *nhr-2*(RNAi) and *nhr-180*(RNAi) treatment lead to faster growth of *atfs-1(null)* animals. **(B)** Bar graphs plotted as % of adults after 55 hrs from egg stage. N =3, biologically independent replicates. \**p*< 0.05 Error bars mean ± SD (one-way ANOVA). **C, D** Representative images of TMRE-stained wildtype, *atfs-1(null),* and *atfs-1(null); nhr-180*(RNAi). Scale bar, 5μm. N =3, biologically independent replicates. **D** Bar graph showing the mean TMRE intensity per area. N=15 animals, \*\*\**p* < 0.0001 (one-way ANOVA). **E-G** Growth as % adults of *clk-1(qm30)* mutants and *isp-1(qm150)* mutants (**F, G**) after control, *nhr-2* or *nhr-180* RNAi from the egg stage. N =3, biologically independent replicates. \**p*< 0.05 Error bars mean ± SD (one-way ANOVA). **H, I** Lifespan graphs of wildtype (N2), *atfs-1(null)* and *atfs-1(null); nhr-180*(RNAi) animals. N =3, biologically independent replicates. \**p* < 0.01 (log-rank test). **J** Quantification of mtDNA by qPCR in *atfs-1(null)* worms raised on control(RNAi) or *nhr-180*(RNAi). N = 3, biologically independent samples. \*\**p* < 0.01 (two-tailed Student’s t-test).

Impressively, inhibition of *nhr-180* increased the developmental rate of *atfs-1(null)* worms to a level comparable to wildtype worms (Fig. 1B, C). Thus, we focused the remainder of our study on *nhr-180*. We next examined the impact of *nhr-180* inhibition on the growth rate of additional mutant strains with dysfunctional mitochondrial, including *clk-1(qm30),* which impairs ubiquinone biosynthesis, and *isp-1(qm150),* which has impairs cytochrome c oxidoreductase activity (OXPHOS complex III) (Felkai et al., 1999; Feng et al., 2001). Interestingly, *nhr-180* inhibition in OXPHOS mutant strain via RNAi increased the developmental rate. 72% of *clk-1(qm30)* and 84% for *isp-1(qm150)* worms developed into adults compared to 5% and 16% in control RNAi after 58hrs of egg laying (Figs. 1E-G). To determine the impact of *nhr-180* inhibition on mitochondrial function, we stained the each strain with TMRE (Tetramethyl rhodamine, Ethyl Ester, Perchlorate), which accumulates within the matrix of functional mitochondria due to the inner membrane potential generated by a functional OXPHOS system (Mallick and Haynes, 2024; Yang et al., 2022). As expected, TMRE staining was reduced in *atfs-1(null)* worms relative to wildtype worms (Fig. 1D). However, TMRE staining is increased in *atfs-1(null)* worms when raised on *nhr-180*(RNAi) to levels comparable to wildtype worms (Fig. 1C-D). Consistent with increased mitochondrial function, *nhr-180*(RNAi) also had increased mitochondrial DNA (mtDNA) content and an extended lifespan relative to *atfs-1(null)* worms (Fig. 1H-J).

We next determined the impact of *nhr-180* inhibition on mitochondrial function in worms with loss-of-function mutations in OXPHOS genes. Impressively, when raised on *nhr-180*(RNAi), *isp-1(qm150)*, *clk-1(qm30)*, or *mev-1(kn1)* (impaired OXPHOS complex II) mutant worms have increased TMRE staining relative to worms raised on control (RNAi) consistent with increased mitochondrial function. Importantly, TMRE staining was also increased in *lonp-1(null)* worms, which lack an essential mitochondrial matrix-localized quality control protease (Supplementary Figs. 1A-F). Combined, these data indicate that *nhr-180* inhibition increases mitochondrial function in worms with impaired respiratory complex components, in worms lacking *atfs-1*, as well as worms with impaired mitochondrial proteostasis. Moreover, these findings indicate that *nhr-180*-dependent transcription contributes to, or exacerbates, the mitochondrial dysfunction observed in *atfs-1(null)* worms as well as in the strains with impaired OXPHOS function.

To further evaluate the impact of *nhr-180* deletion on mitochondrial function during stress, we introduced 4 stop codons in the first exon of the *nhr-180* gene that encodes the DNA binding domain via CRISPR-Cas9-mediated genome editing (Fig. 2A). Consistent with our *nhr-180*(RNAi) results (Fig. 1I), *atfs-1(null);nhr-180(cmh19)* deletion worms had a significantly increased lifespan relative to *atfs-1(null)* worms (Fig. 2B-D). Lastly, *atfs-1(null);nhr-180(cmh19)* worms also had an increased body bending and thrashing rate compared to *atfs-1(null)* worms (Supplementary Fig. 2A-B) consistent with increased mitochondrial function. Similar to the phenotypes observed when raised on *nhr-180*(RNAi), *atfs-1(null);nhr-180(cmh19)* worms also had increased TMRE staining, mtDNA content, and respiration compared to *atfs-1(null)* worms which is consistent with a more robust mitochondrial network upon *nhr-180* inhibition (Fig. 2E-G, 2I-J, Supplementary Fig. 2C-D). These findings suggest that the improved fitness of *atfs-1(null)* worms that occurs upon reduced NHR-180 expression is due to increased mitochondrial function.

**Figure 2:**
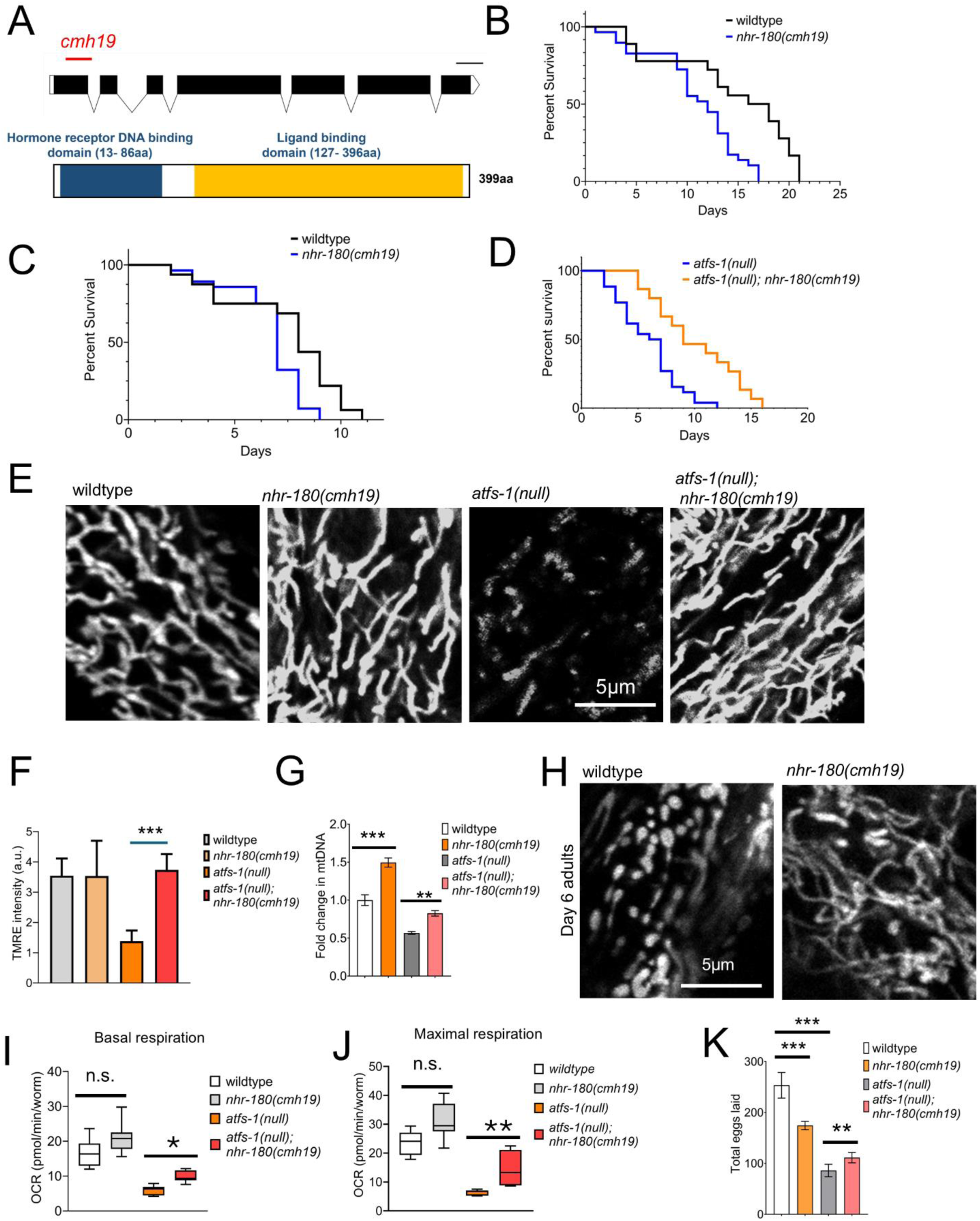
*nhr-180(cmh19)* mutant worms show increased TMRE staining and mtDNA content. **A** schematic showing the exons (boxes) and introns (lines) of the NHR-180 protein with the conserved domains. *cmh19* CRISPR allele with multiple stop codons is shown in red. **B, C** Lifespan graphs for wildtype and *nhr-180* mutants at 20°C (**B**) and 25°C (**C**). **D** Lifespan graph for *atfs-1(null)* and *atfs-1(null); nhr-180(cmh19)* mutants at 20°C. N =3, biologically independent replicates. \**p* < 0.01 (**B, D**) n.s. (**C**) (log-rank test). **E** Representative images of TMRE stained wildtype, *nhr-180(cmh19), atfs-1(null)* and *atfs-1(null); nhr-180(cmh19)* at the young adult stage. Scale bar, 5μm. N =3, biologically independent replicates. **F** Bar graph showing the mean TMRE intensity per area. N=15 animals per condition, \*\*\**p* < 0.0001 (one-way ANOVA). **G** Quantification of mtDNA by qPCR in wildtype, *nhr-180(cmh19), atfs-1(null)* and *atfs-1(null); nhr-180(cmh19)*. N = 3, biologically independent samples. \*\**p* < 0.01 (one-way ANOVA). **H** Representative images of TMRE stained wildtype and *nhr-180(cmh19)* at day-6 of adult stage. Scale bar, 5μm. N =3, biologically independent replicates. **I, J** Bos and violin plot showing the oxygen consumption in wildtype, *nhr-180(cmh19), atfs-1(null)* and *atfs-1(null); nhr-180(cmh19)* worms at the L4 stage. N = 3, biologically independent samples. \**p* < 0.05, \*\**p* < 0.01 (multiple t-test). **J** Bar graphs showing the total number of eggs laid in wildtype, *nhr-180(cmh19), atfs-1(null)* and *atfs-1(null); nhr-180(cmh19)* worms N = 3, biologically independent samples. \*\**p* < 0.01, \*\*\**p* < 0.001 (one-way ANOVA).

In addition to a decline in mitochondrial function, fecundity is also known to decline as worms age (Gaffney et al., 2018). Intriguingly, we found that at adult day 4, *nhr-180(cmh19)* worms have significantly more TMRE staining relative to wildtype worms (Fig. 2H). While *nhr-180* mutants have a reduced brood size compared to wildtype worms, *atfs-1(null);nhr-180(cmh19)* worms laid more eggs than *atfs-1(null)* worms (Fig. 2K), consistent with increased health span. Combined, our data suggest that impaired expression of NHR-180 promotes mitochondrial function in worms with severe mitochondrial dysfunction, caused by perturbations within OXPHOS components or UPR^mt^ inhibition.

### NHR-180 is expressed in intestinal, hypodermal, and muscle cells

To identify the cells or tissues in which NHR-180 is expressed, we generated a translational reporter where NHR-180 was tagged with GFP at the C-terminus (*nhr-180_pr_::nhr-180::GFP*) (Fig. 3A). We found that NHR-180::GFP is expressed in eggs and in several tissues throughout development and into adulthood. While NHR-180::GFP expression is most obvious in the intestine, NHR-180::GFP is also expressed in muscle cells as well as hypodermal cells (Fig. 3B). Within intestinal cells, NHR-180::GFP is located in both the cytosol and the nucleus. However, in muscle and hypodermal cells, NHR-180::GFP is primarily in the nucleus (Fig. 3B). To confirm that the GFP expression correlates with the levels of NHR-180, we observed a reduction in GFP fluorescence when worms were raised on *nhr-180*(RNAi) (Fig. 3E).

**Figure 3:**
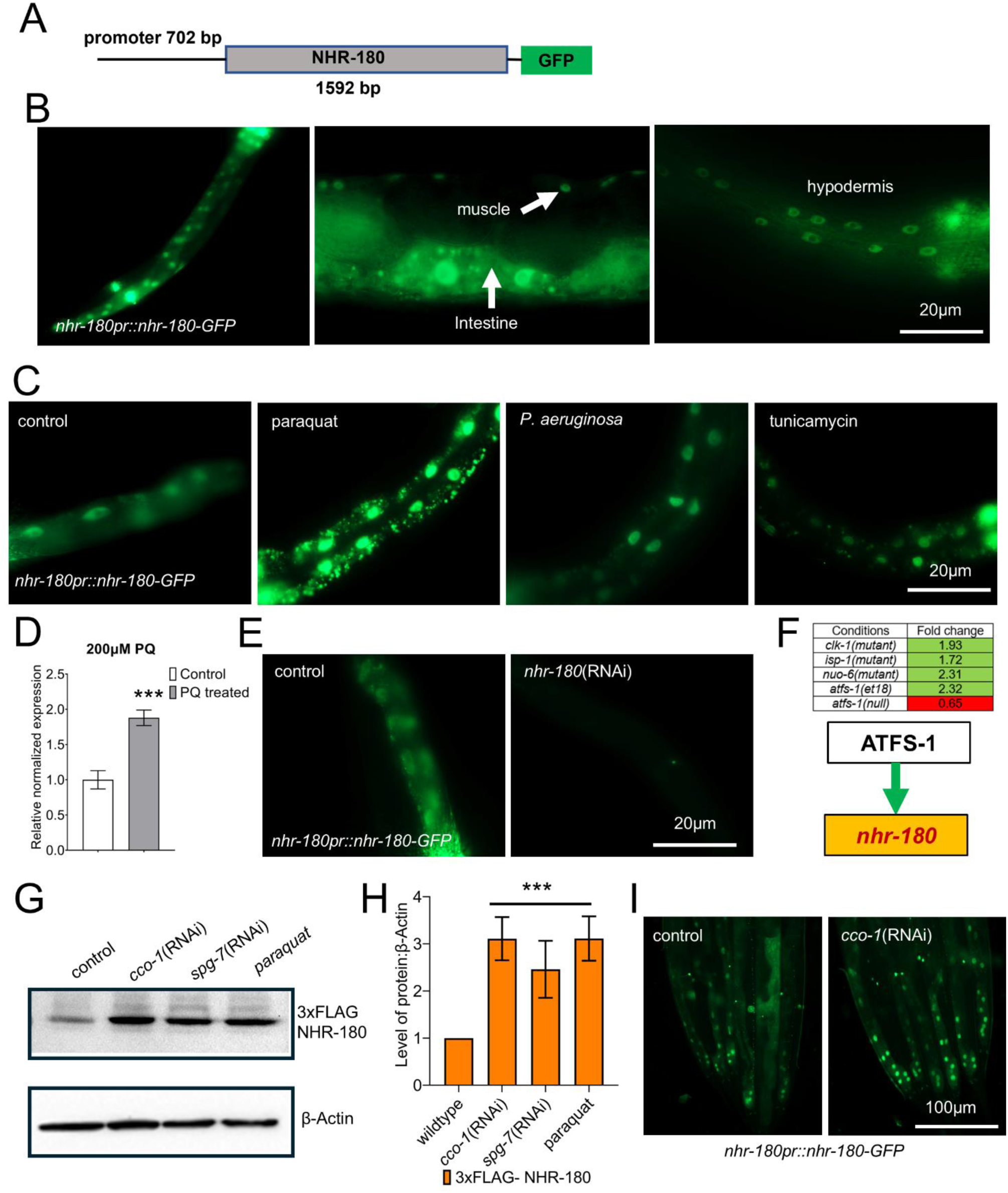
NHR-180 is expressed in the intestine, muscle, and hypodermis. **A** Schematic showing the plasmid construct of *nhr-180_pr_::NHR-180-GFP* translational reporter. **B** Representative images of the NHR-180-GFP strain in the intestinal cells, muscle cells, and hypodermal cells. Scale bar, 20μm **C** Representative NHR-180-GFP images at the young adult stage after growing worms from the L4 stage on untreated *E. coli* plates (control), *E. coli* plates treated with paraquat, *P. aeruginosa* plates, and *E. coli* plates treated with tunicamycin. Scale bar, 20μm. **D** Quantification of *nhr-180* transcripts by qPCR in wildtype worms with and without paraquat treatment. N = 3, biologically independent samples. \*\*\**p* < 0.001 (two-tailed Student’s t-test). **E** Representative NHR-180-GFP images at the young adult stage after control(RNAi) and *nhr-180(*RNAi). Scale bar, 20μm. **F** Table showing the fold change in the *nhr-180* expression in different mitochondrial mutants. **G** Western blot showing 3xFLAG::NHR-180 and β-Actin. **H** Bar graph showing the quantification of the western blot in **G**. **H** Representative NHR-180-GFP images at the young adult stage following control or *cco-1*(RNAi) bacteria, scale bar represents 100µm.

We next sought to determine the physiologic or pathologic conditions that induce NHR-180 expression. We exposed *nhr-180_pr_*::*nhr-180*::gfp worms to paraquat (oxidative stress), *P. aeruginosa* (infection), or tunicamycin (ER stress). Interestingly, only paraquat treatment increased the expression of NHR-180::GFP, which accumulated within the cytosol and the nucleus (Fig. 3C). Additionally, *nhr-180* transcripts are also increased in wildtype worms upon paraquat treatment as determined by qRT-PCR (Fig. 3D), consistent with paraquat causing mitochondrial dysfunction (Bora et al., 2021; Runkel et al., 2013). These findings are consistent with previously published transcriptomic data showing that *nhr-180* transcripts are induced upon exposure to paraquat in wildtype worms, but also induced in mutant worms with impaired OXPHOS, including *isp-1(qm150)*, *clk-1(qm30)*, or *nuo-6(qm200)* worms (Felkai et al., 1999; Yang and Hekimi, 2010). Mutations in several OXPHOS components *isp-1(qm150)*, *clk-1(qm30), nuo-6(qm200)* induce *atfs-1*-dependent transcription, resulting in increased *nhr-180* transcription (Fig. 3F). Furthermore, mutations that perturb the ATFS-1 mitochondrial targeting sequence and activate *atfs-1*-dependent transcription also induce transcription of *nhr-*180 (Rauthan et al., 2013; Shpilka et al., 2021). Next, we sought to determine if the NHR-180 protein levels increase after mitochondrial perturbations that promote nuclear localization of ATFS-1. For these studies, we introduced a 3xFLAG epitope at the amino-terminus of NHR-180 via CRISPR-Cas9 genome editing. We found that NHR-180 protein levels increased upon OXPHOS inhibition, *cco-1*(RNAi), mitochondrial protease inhibition – *spg-7*(RNAi), and exposure to paraquat, which generates excessive reactive oxygen species (Fig. 3G and H). Combined, these data indicate that NHR-180 expression is increased in response to a variety of mitochondrial perturbations.

### *nhr-180* regulates transcription of genes required for protein synthesis in the cytosol and mitochondria

To gain insight into the role of NHR-180 in regulating mitochondrial function, we carried out transcriptomic analysis comparing wildtype, *nhr-180(cmh19)*, *atfs-1(null)*, and *atfs-1(null);nhr-180(cmh19)* worms at the L4 stage. We found that ∼1380 genes are upregulated in *nhr-180(cmh19)* worms and ∼750 genes are downregulated compared to wildtype worms (Fig. 4A and B, Supplementary Table 2). Interestingly, gene enrichment analyses indicated that genes required for cytosolic protein synthesis as well as protein synthesis within mitochondria on mitochondrial ribosomes are reduced in *nhr-180(cmh19)* worms (Fig. 4C, Supplementary Tables 3 and 4).

**Figure 4:**
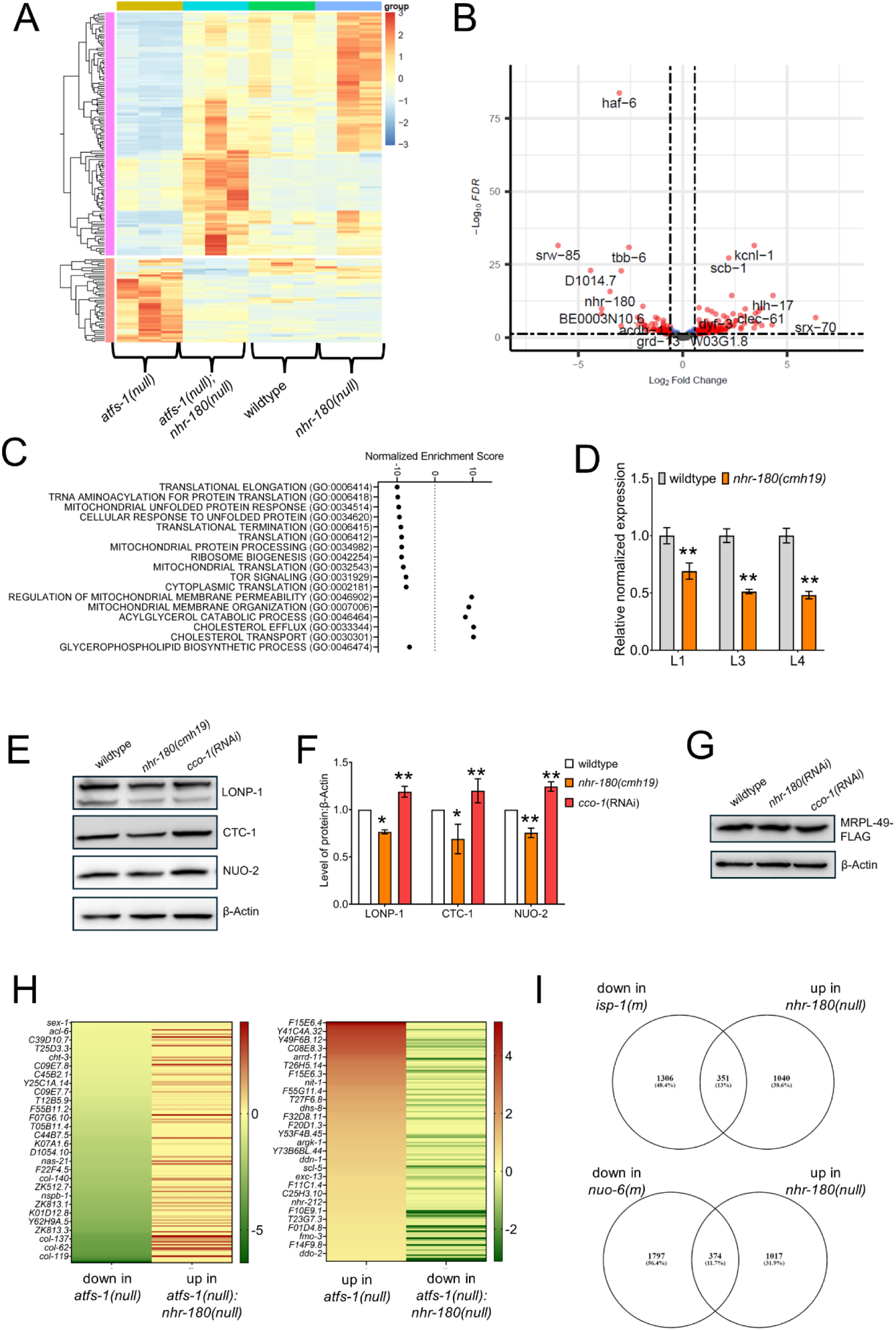
Loss of NHR-180 reverses gene expression in *atfs-1(null)* and other OXPHOS mutants. **A** Heatmap showing gene expression changes in wildtype (N2), *nhr-180(null), atfs-1(null)* and *atfs-1(null); nhr-180(null)* mutants. **B** MA plot showing differentially expressed genes in *nhr-180(cmh19)* mutants versus wildtype (N2). **C** Bubble plot showing the normalized enrichment scores of gene ontology classes for differentially expressed genes in the *nhr-180* mutant. **D** Quantification of UPR^mt^ reporter *hsp-6* transcripts by qPCR in wildtype, and *nhr-180(cmh19)* at different larval stages. N = 3, biologically independent samples. \*\*\**p* < 0.001 (one-way ANOVA). **E** Western blot showing LONP-1, CTC-1, NUO-2, and β-Actin in wildtype, *nhr-180(cmh19),* and *cco-1*(RNAi) treated worms. **F** Bar graph showing the quantification of western blot in E. \**p* < 0.05 and \*\**p* < 0.01 (one-way ANOVA). **G** Western blot showing MRPL-49::3xFLAG in wildtype, *nhr-180(cmh19),* and *cco-1*(RNAi) treated worms. **H** Heatmap listing the genes in *atfs-1(null)* that are reversed in the double mutants of *atfs-1(null); nhr-180(cmh19).* **I** Venn diagrams showing the common sets of genes between OXPHOS mutants and *nhr-180* mutants.

Eukaryotic initiation factor 3 complex components *eif-3.B, eif-3.d, eif-3e, eif-3h,* and *eif-3.l*) are also reduced in *nhr-180(cmh19)* worms. Interestingly, inhibition of any of the eIF3 components that are downregulated in *nhr-180(cmh19)* worms increased the development rate in *atfs-1(null)* worms, similar to that seen upon *nhr-180* inhibition (Supplementary Fig. 3, Supplementary Table 3). Intriguingly, genes induced by ATFS-1, such as *hsp-6* and *lonp-1,* are also reduced in *nhr-180(cmh19)* worms, suggesting that NHR-180 may contribute to mitochondrial stress response (Fig. 4C and D). Consistent with our RNAseq data, the mitochondria matrix-localized protease LONP-1, as well as the OXPHOS proteins CTC-1 (complex I) and NUO-2 (OXPHOS complex I component), are reduced in *nhr-180(cmh19)* worms (Fig. 4E-G). Moreover, 568 genes that are highly expressed during larval development and L4 stage are upregulated in *nhr-180(cmh19)* (Fig. 4H and I, Supplementary Table 5), suggesting that increased *nhr-180* expression slows larval development, a phenotype observed in the OXPHOS mutants and *atfs-1(null)* worms (Hendriks et al., 2014; Jackson et al., 2014). Combined, our data suggest that *nhr-180* regulates a transcription program that promotes protein synthesis in both the mitochondrial matrix and on cytosolic ribosomes. Furthermore, *nhr-180*-dependent transcription suppresses proteolytic genes such as *ubc-24*, *marc-2*, *fbxa-125*, *fbxa-183*, and *fbxa-224*, which are also repressed in OXPHOS mutants such as *isp-1(qm150)* and *nuo-6(qm200)* worms (Yee et al., 2014).

### AMP binds the ligand-binding domain of NHR-180

We next sought to identify the mechanism, or ligand, that stimulates *nhr-180*-dependent transcription that occurs during mitochondrial dysfunction caused by paraquat, inhibition of complex III component *isp-1*, ubiquinone biosynthesis component *clk-1* or impaired expression of *cco-1*, or *spg-7* via RNAi. Because OXPHOS inhibition via impaired expression of *cyc-1*, *nuo-2,* and *cco-1*, is known to cause a reduction in ATP (Dillin et al., 2002) we hypothesized that increased AMP accumulation binds the ligand-binding domain within NHR-180 and increases *nhr-180*-dependent transcription. To determine if AMP can bind the ligand-binding domain of NHR-180, we used AlphaFold prediction software (Abramson et al., 2024). Interestingly, AlphaFold predicted that AMP interacts with the ligand-binding domain of NHR-180 via five hydrogen bonds (Fig. 5A-D). To disrupt the predicted AMP binding site within NHR-180, we mutated the three predicted hydrogen-bonding amino acids (E180, N184, and W185) to (I, L, and P) (Fig. 5E). We then compared the expression of the mitochondrial chaperone *hsp-6,* ATP citrate synthase *acly-1,* cytosolic translation elongation factors*-eif-3c and eif-3l,* and mitochondrial translation elongation factors*-tufm-1*, and *tsfm-1* mRNAs, in wildtype, *nhr-180(null)*, and *nhr-180^I,L,P^* worms. Importantly, all of the tested mRNAs were reduced in *nhr-180(null)* and *nhr-180^I,L,P^* worms compared to wildtype worms (Fig. 5F and G). Combined, these findings indicate that these three amino acids are required for NHR-180 regulation.

**Figure 5:**
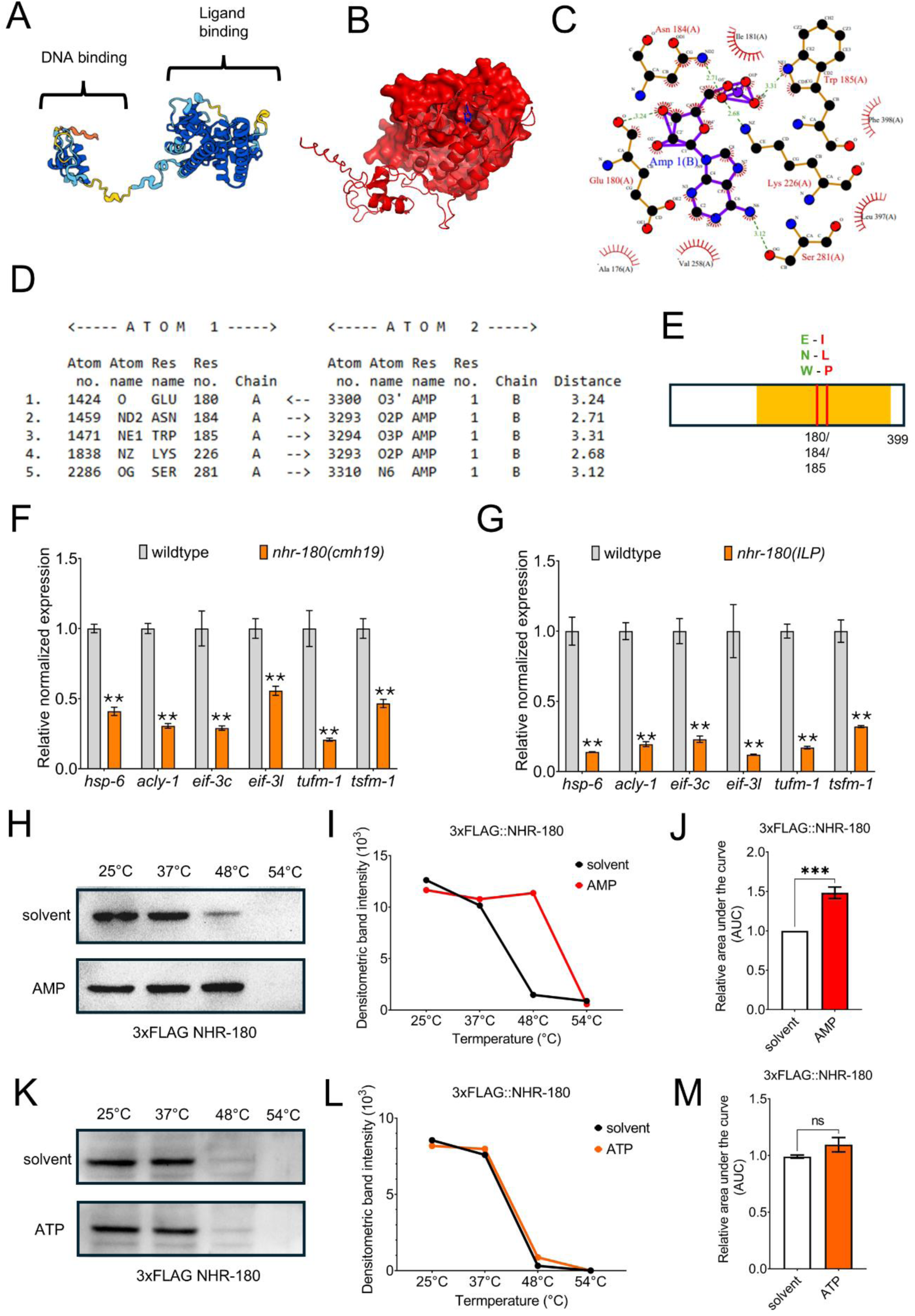
AMP is predicted to bind at the ligand-binding domain of NHR-180. **A** Secondary structure of full-length NHR-180 showing the DNA- and ligand-binding domains. **B** *In silico* molecular modeling of full-length NHR-180 (red). The identified ligand-binding pocket for AMP is indicated in blue. **C** *In silico* model of intramolecular interactions (hydrogen bonding) between the ligand binding domain of NHR-180 and AMP. **D** Table showing the list of five potential hydrogen bond interactions between the amino acids in the ligand binding domain of NHR-180 and AMP. **E** Schematic diagram of NHR-180 protein showing the position of three hydrogen-bonded amino acid residues (E, N, and W), CRISPR mutated to I, L, and P. **F-G** qPCR analysis of the target genes (*hsp-6* – mitochondrial chaperone, *acly-1* – ATP citrate synthase, *eif-3.c/l* – translation initiation factor 3 complex components, *tufm-1* – mitochondrial translation elongation factor, and *tsfm-1* - mitochondrial translation elongation factor) of NHR-180 in wildtype, *nhr-180(cmh19),* and *nhr-180(ILP)* at L4 stage. Data represents the mean of biological replicates (N = 3) with error bars indicating the standard error of the mean (SEM). \*\**p* < 0.01 (one-way ANOVA). **H, K** A representative immunoblot of a cellular thermal shift assay (CETSA) experiment using an anti-FLAG antibody that probed whole-cell lysates from a transgenic *C. elegans* strain in which NHR-180 was tagged with 3xFLAG at its endogenous locus. **I, L** A representative densitometric quantification from a CETSA experiment that characterized the interaction of AMP (100µM), ATP (100µM), and solvent (water) with 3xFLAG::NHR-180. **J, M** The area under the curve was quantified from each biological replicate for the experiment described in **(I)** and **(L)** and normalized to the solvent control (n=3). Data in J and M represent the average of all biological replicates, with error bars indicating the standard error of the mean (SEM). *** *p* < 0.001 (two-tailed, unpaired t-test).

As an orthologous means to determine if AMP binds NHR-180, we utilized a cellular thermal shift assay (CETSA), a technique based on the principle that the binding of a ligand to the NHR ligand-binding domain stabilizes the domain and protects the protein complex against denaturation and aggregation upon exposure to heat (Jafari et al., 2014; Tse and Pukkila-Worley, 2023). Using the 3xFLAG-NHR-180 strain, we probed for NHR-180^FLAG^ in whole-cell lysates using an anti-FLAG antibody. Interestingly, incubation with AMP promoted NHR-180 stabilization over a range of temperatures (Fig. 5H). We quantified the area under the curve from biological replicates and found that treatment with AMP increased the thermal stability of NHR-180 (Fig. 5I and J). Importantly, unlike treatment with AMP, incubation with ATP did not stabilize NHR-180, suggesting that AMP is the activating ligand that stimulates NHR-180-dependent transcription (Fig. 5K-M). As a control, we used two strains expressing 3xFLAG-labelled NHR-12 and NHR-86 proteins. AMP failed to stabilize either NHR-12 or NHR-86 (Supplementary Fig. 4A-D). These results indicate that AMP binding promotes NHR-180-dependent transcription.

### Inhibition of protein synthesis increases mitochondrial function

Caloric restriction leads to AMPK activation and the downstream phosphorylation of S6 kinase, which leads to reduced protein synthesis (Burkewitz et al., 2014; Salminen and Kaarniranta, 2012). We compared the transcription profiles of *eat-2(ad465)* worms, which are calorie-restricted due to a defect in pharyngeal pumping, to *nhr-180(cmh19)* (Lakowski and Hekimi, 1998). Interestingly, 554 of the upregulated genes in *nhr-180* mutants are also upregulated in the *eat-2* mutants (Fig. 6A, Supplementary Table 6) (Heestand et al., 2013). We next compared the transcriptional profile of *nhr-180(cmh19)* worms with a strain that expresses the constitutively active allele of AMPK (CA-AAK-2), which leads to extended lifespan and improved mitochondrial function (Weir et al., 2017). Genes upregulated in *nhr-180(cmh19)* worms significantly overlapped with those upregulated in the CA-AAK-2 mutant as well (Supplementary Table 7). (Chen et al., 2013; Mair et al., 2011).

**Figure 6:**
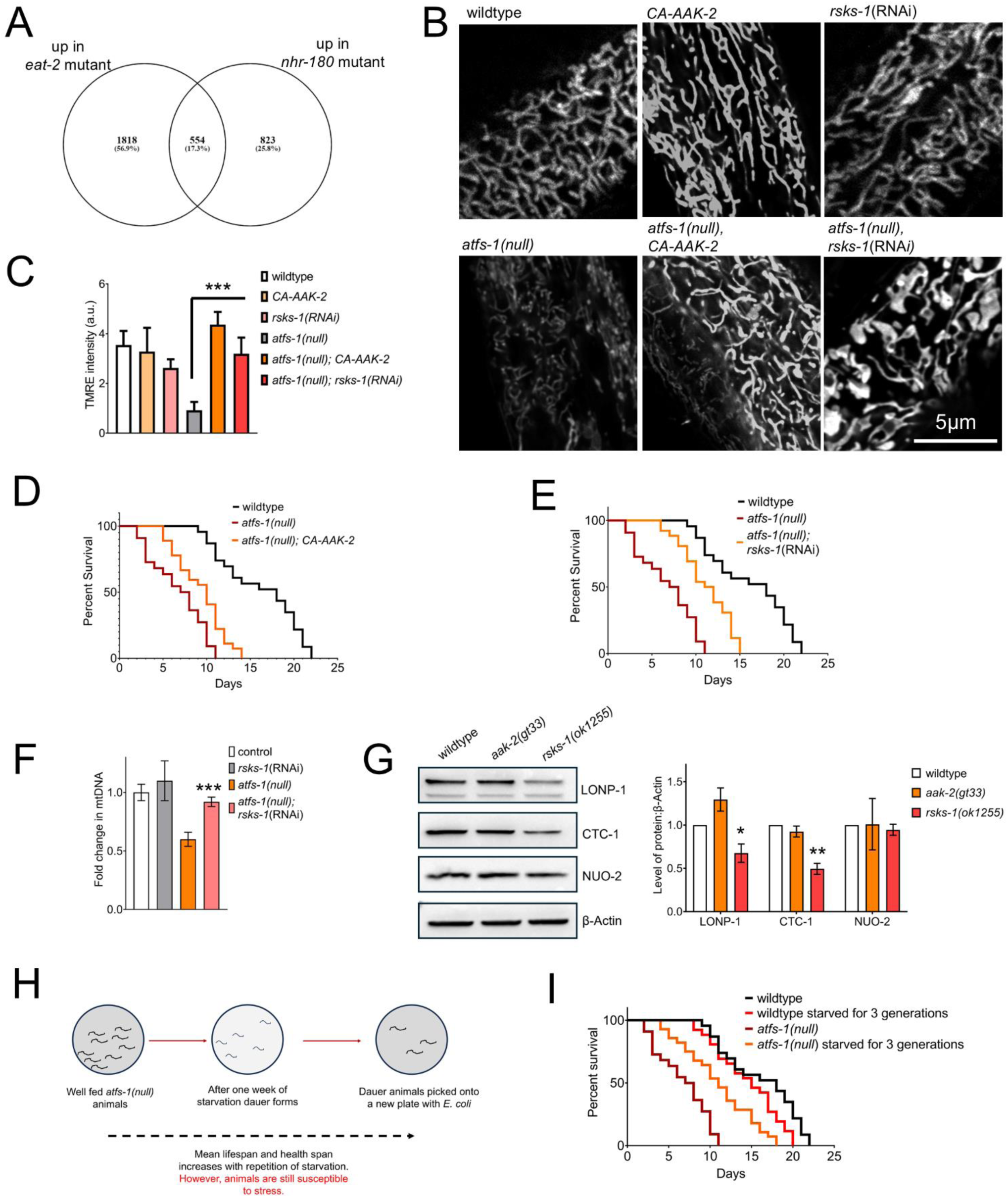
Activating AAK-2/AMPK or inhibiting RSKS-1/S6K promotes mitochondrial function. **A** Venn diagram shows the common sets of genes between *eat-2* mutant and *nhr-180* mutants. (See supplementary Table 6 for a complete list of genes). **B** Representative images of TMRE stained wildtype, CA-AAK-2, *rsks-1*(RNAi), *atfs-1(null)*, *atfs-1(null); CA-AAK-2* and *atfs-1(null); rsks-1*(RNAi) at the young adult stage. Scale bar, 50μm. **C** Bar graph showing the mean TMRE intensity per area. Data presents average of 3 biological replicates and error bars as SEM, with N=15 animals per condition, \*\*\**p* < 0.0001 (one-way ANOVA). **D, E** Lifespan graphs of wildtype (N2), CA-AAK-2, *rsks-1*(RNAi), *atfs-1(null)*, *atfs-1(null); CA-AAK-2* and *atfs-1(null); rsks-1*(RNAi) animals. N =3, biologically independent replicates. \**p* < 0.01 (log-rank test). **F** qPCR analysis of total mtDNA in wildtype and *atfs-1(null)* following control(RNAi) or *rsks-1*(RNAi). N = 3, biologically independent samples. \*\**p* < 0.01 (one-way ANOVA). **G** A representative immunoblots for endogenous LONP-1, CTC-1, NUO-2 and β-Actin in wildtype, *aak-2(gt33)* and *rsks-1(ok1255)* adult worms. The bar graph represents the average of 3 biological replicates of immunoblot band intensities with error bars giving SEM. * *p* < 0.05 and ** *p* < 0.01 (two-tailed t-test). **H** Schematic showing the starvation steps followed for wildtype (N2) and *atfs-1(null)* worms that improved lifespan (**H**) and other phenotypes. **I** Lifespan graphs of wildtype and *atfs-1(null)* worms, either continuously fed on food or intermittent starvation between generations. Statistics are done using 3 biological replicates with more than 50 animals per condition, *p* < 0.01 using the log-rank test.

We next examined if the constitutively active form of the AMPK catalytic subunit AAK-2 (CA-AAK-2) or inhibition of the S6 kinase homolog RSKS-1 affected mitochondrial function and the development of *atfs-1(null)* worms, as both reduce the rate of protein synthesis (Burkewitz et al., 2014; Herzig and Shaw, 2018). Impressively, expression of CA-AAK-2 or inhibition of *rsks-1* expression increased TMRE staining in *atfs-1(null)* worms to a level similar to that observed in wildtype worms (Fig. 6B and C). Importantly, the lifespan of *atfs-1(null)* worms also increased significantly (Fig. 6D and E). As an alternative approach to evaluating mitochondrial biogenesis and function, we determined the mtDNA content in the *atfs-1(null)* worms when raised on *rsks-1*(RNAi). Consistent with the increased mitochondrial function of *atfs-1(null)* worms raised on *rsks-1*(RNAi) (Fig 6B and C), mtDNA content was also increased as determined by qRT-PCR (Fig. 6F). Furthermore, the levels of LONP-1, CTC-1 and NUO-2 were reduced in the *rsks-1(ok1255)* worms supporting the role of RSKS-1/S6 kinase in promoting protein translation (Fig. 6G). Combined, the data indicate that a reduction in cytosolic protein synthesis increases mitochondrial function in worms with dysfunctional OXPHOS.

We next determined whether starvation also increases mitochondrial function in *atfs-1(null*) worms. We starved *atfs-1(null)* worms until they entered the dauer diapause phase and then re-fed the worms to complete the reproductive cycle. Lifespan and health span were examined after worms were recovered from the dauer phase and fed a bacterial diet (Fig. 6H). Importantly, starved *atfs-1(null)* worms lived significantly longer than continuously fed *atfs-1(null)* worms (Fig. 6I). Moreover, starved *atfs-1(null)* worms had increased TMRE staining and thrashing rates relative to fed *atfs-1(null)* worms (Supplementary Figs. 5A-B). The increased lifespan in *atfs-1(null)* worms was abolished when the worms were refed for two successive generations, suggesting that the phenotype was not due to permanent genetic modification. To determine if AMP kinase is required for the increased mitochondrial function observed in *atfs-1(null)* worms following starvation, we generated *atfs-1(null);aak-2(gt33)* worms, which also shortened lifespan relative to *atfs-1(null)* worms and were unaffected by starvation (Supplementary Fig. 5C-D). These findings indicate that *aak-2/AMPK* is required for the increased mitochondrial function observed in the *atfs-1(null)* worms following starvation.

### Inhibition of S6K1 increases mitochondrial function in OXPHOS-deficient cells

To determine if the impact of S6 kinase inhibition on mitochondrial function is conserved in mammals, we generated ATF5-/- knockout HEK293T cells, which inhibit the mammalian ortholog of ATFS-1 (Fiorese et al., 2016). Similar to *atfs-1(null)* worms, *ATF5*-/- cells have reduced mtDNA content and impaired respiration (Fiorese et al., 2016; Yang et al., 2022). Interestingly, treatment with the S6 kinase-1 inhibitor (PF-4708671) increased mtDNA content, as well as basal and maximal respiration in *ATF5*-/- cells, suggesting that inhibition of S6 kinase promotes mitochondrial function in mammalian cells (Fig. 7A-E) (Jiang et al., 2022). We also examined a heteroplasmic cybrid cell line that harbors a combination of wildtype mtDNAs and mutant mtDNAs harboring a 4,977-base-pair mtDNA deletion, known as the ‘common deletion’(King and Attardi, 1989), which removes multiple OXPHOS genes. Accumulation of the deleterious genome is associated with Kearns–Sayre syndrome (KSS), progressive external ophthalmoplegia, cancer, and ageing (Moraes et al., 1989). Interestingly, we found that the inhibition of S6K1 via the inhibitor (PF-4708671) or siRNA increased respiration and TMRE staining in the KSS cybrid cells (Fig. 7F-G and 7I-J). Lastly, we examined the impact of PF-4708671 on a cell line with impaired succinate dehydrogenase complex (II) caused by a gene deletion (Spinelli et al., 2021). Upon exposure to PF-4708671, basal respiration increased (Fig. 7H). Together, these findings indicate that reduced cytosolic protein synthesis via S6 kinase inhibition enhances mitochondrial function to a level similar to wildtype cells.

**Figure 7:**
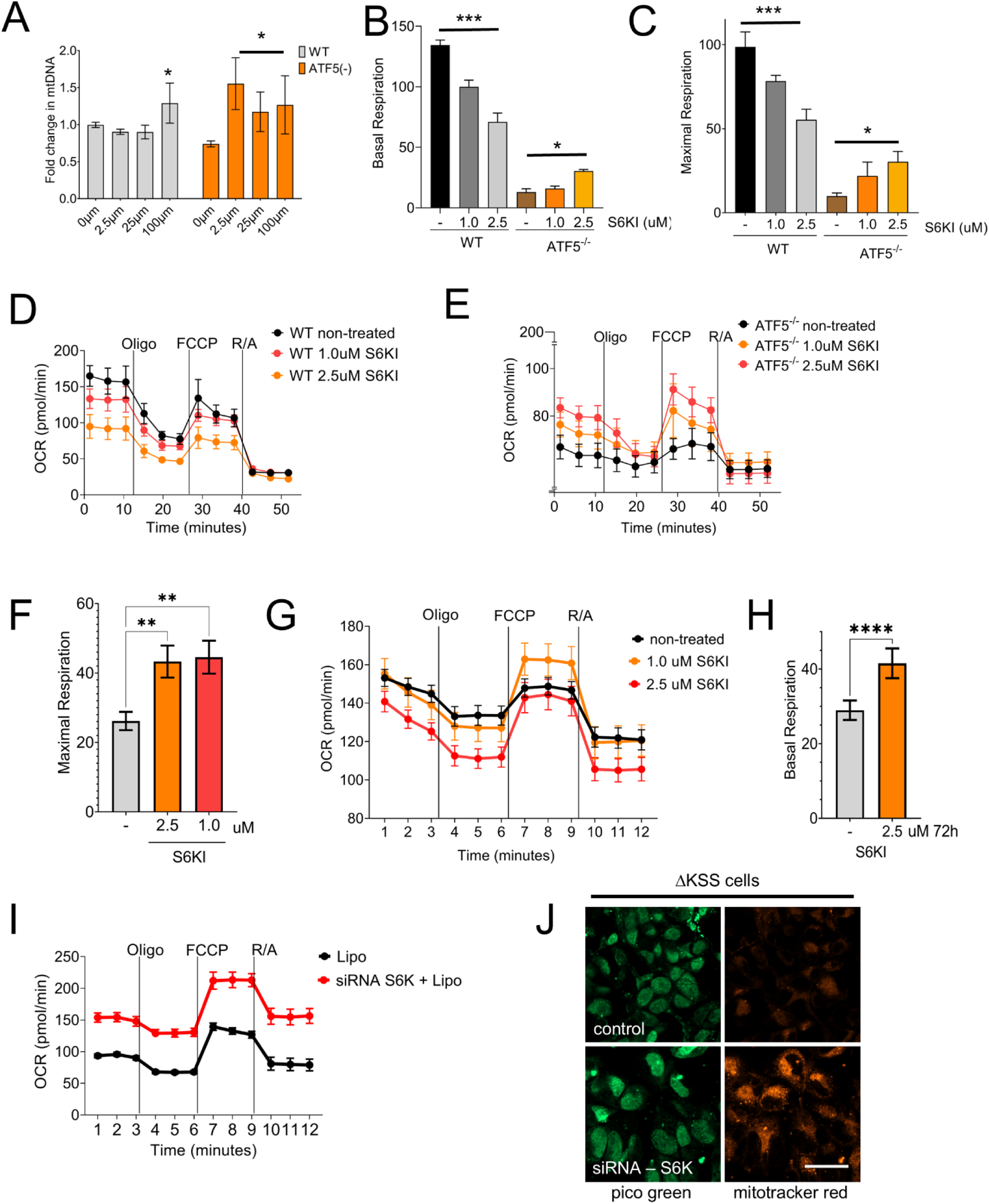
Inhibiting S6 kinase improves oxygen consumption in OXPHOS-deficient cells. **A** qPCR analysis of total mtDNA content in HEK293T cells in control and ATF5 KO treated with DMSO, 2.5µm, 25µm, and 100µm of PF-4708671 for 48 hrs. N = 3, biologically independent samples. \*\**p* < 0.01 (one-way ANOVA). **B, C** Quantification of basal and maximal respiration in control and ATF5 KO HEK293T cells treated with DMSO, 1µm, and 2.5µm of PF-4708671 for 48 hrs. **A-C** Biologically 3 independent samples, * *p* < 0.05 and *** *p* < 0.001 (one-way ANOVA). Bar graphs represent the average of the replicates, with error bars indicating the SEM. **D, E** Oxygen consumption rate (OCR - pmol/min) in HEK293T cells in control and ATF5 KO treated with DMSO, 1µm, and 2.5µm of PF-4708671 for 48 hrs. **F** Quantification of maximal respiration in ΔKSS cybrid cells treated with DMSO, 1µm, and 2.5µm of PF-4708671 for 48 hrs. **G** OCR in ΔKSS cybrid cells treated with DMSO, 1µm, and 2.5µm of PF-4708671 for 48 hrs. **H** Quantification of maximal respiration in SDHA KO cells treated with DMSO and 2.5µm of PF-4708671 for 72 hrs. **F, H** Biologically 3 independent samples, ** *p* < 0.01 and **** *p* < 0.0001 (one-way ANOVA). Bar graphs represent the average of the replicates, with error bars indicating the SEM. **I** OCR in ΔKSS cybrid cells treated with or without S6 kinase siRNA. **J** Representative images of ΔKSS cells stained with Pico green (stains double-stranded DNA) and Mito tracker red (stains functional mitochondria) after treating with empty vector or S6K siRNA. The scale bar represents 10µm. Three biological replicates with similar outcomes.

## Discussion

The UPR^mt^ was discovered as a pathway that responds to mitochondrial perturbations, including impaired mitochondrial protein import or impaired OXPHOS (Durieux et al., 2011; Nargund et al., 2012; Shpilka et al., 2021). Here, we identify a previously unstudied transcription factor that is also activated upon mitochondrial stress or dysfunction. Mitochondrial dysfunction impedes ATP synthesis, which increases AMP levels. Here, we demonstrate that AMP binds NHR-180 and promotes NHR-180-dependent transcription. In *nhr-180* mutant worms, mitochondrial protein transporters, mitochondrial translation elongation factors, mitochondrial ribosomal subunits, and cytosolic translation initiation factors are reduced, suggesting that NHR-180 promotes transcription of genes required for protein synthesis both in the cytosol and the mitochondria. Interestingly, we demonstrate that by inhibiting *nhr-180* expression, mitochondrial function can be increased in *atfs-1*-deletion worms or worms with mutations that impair OXPHOS. We propose a model whereby cells respond to transient mitochondrial dysfunction by activating *nhr-180,* which promotes protein synthesis on cytosolic ribosomes as well as on mitochondrial ribosomes. We found that NHR-180 promotes mitochondrial protein synthesis while concurrently slowing worm development, suggesting a trade-off where enhancing mitochondrial biogenesis takes precedence over rapid development. This shift may represent an adaptive strategy, prioritizing mitochondrial functions to support survival under stress. Such a mechanism highlights the evolutionary importance of balancing energy production with developmental timing in response to mitochondrial dysfunction.

An unbiased screen in yeast cells previously identified 40 suppressors of mitochondrial dysfunction-induced cell death (Haynes, 2015; Wang and Chen, 2015). These suppressors included five classes of genes associated with the target of rapamycin (TOR) signaling, mRNA silencing and turnover, ribosomal function, protein synthesis, tRNA methylation, and cytosolic protein degradation and chaperone expression (Wang and Chen, 2015). A separate study analyzed all of the RNA transcripts and proteins altered in cells in which mitochondrial import is impaired. Notably, the expression and production of many genes and proteins required for protein synthesis were reduced, as was overall protein synthesis (Wrobel et al., 2015). Furthermore, mTOR inhibition that slows growth and reduces protein translation has been shown to mitigate mitochondrial dysfunction-associated phenotypes in mice (Johnson et al., 2013). These independent findings align conceptually with the beneficial effects of inhibiting *nhr-180* observed in worms, where reduced cytosolic and mitochondrial protein synthesis promotes mitochondrial function.

Glucose is required for OXPHOS to generate ATP. However, excessive glucose can perturb mitochondrial function due to reactive oxygen species (ROS) generation inside mitochondria (Allen et al., 2005; Robertson et al., 2003). Intriguingly, research in *C. elegans* shows that impaired glucose metabolism leads to increased mitochondrial respiration and prolonged lifespan (Schulz et al., 2007), which requires AMPK (Cantó and Auwerx, 2011; Herzig and Shaw, 2018). Similarly, dietary restriction or activated AMPK delays the age-associated decline in mitochondrial function in both worms and mice (Lanza et al., 2012; Weir et al., 2017). Our findings that recovery of mitochondrial defects via a reduction of protein synthesis is in line with the effect of dietary restriction on mitochondrial function (Weir et al., 2017), which appears to be conserved (Herzig and Shaw, 2018; Lanza et al., 2012).

In summary, our findings suggest that *C. elegans* evolved a strategy to promote mitochondrial biogenesis during transient mitochondrial dysfunction via the nuclear hormone receptor NHR-180. An increase in AMP increases *nhr-180*-dependent transcription, which includes numerous genes required for translation on cytosolic ribosomes as well as on mitochondrial ribosomes to promote mitochondrial biogenesis and ATP synthesis. We propose that this strategy is advantageous so long as the mitochondrial stress, or dysfunction, is transient. However, if the mitochondrial stress is caused by a mutation within a gene encoding OXPHOS proteins, mitochondrial chaperones, mitochondrial ribosomes, etc., the increase in protein synthesis rate becomes toxic as the dysfunctional mitochondria are unable to process or assemble the imported proteins. Our findings demonstrate that inhibiting cytosolic protein synthesis increases mitochondrial function in OXPHOS-deficient cells. Together, we show a straightforward yet effective strategy to ameliorate mitochondrial dysfunction in disease models caused by mutations in mitochondrial components.

## Materials and Methods

### Worms, plasmids, and staining

N2(wild type), *isp-1(qm150)*, *clk-1(qm30)*, *rsks-1(ok1255),* and *aak-2(gt33)* were obtained from the *Caenorhabditis* Genetics Center (Minneapolis, MN). *nhr-180(cmh19)* and *nhr-180-3x-FLAG* strains were generated via CRISPR-Cas9 in wildtype worms as described (Deng et al., 2019). The crRNAs (IDT) were co-injected with purified Cas9 protein, tracrRNA (Dharmacon), repair templates (IDT), and the *pRF4::rol-6(su1006)* plasmid as described (Dokshin et al., 2018). The *pRF4::rol-6 (su1006)* plasmid was a gift from Craig Mello (Dokshin et al., 2018). *The nhr-180_pr_::nhr-180-GFP* plasmid was generated using conventional cloning of the *nhr-180* coding sequence with 5’ and 3’ UTRs (2241 bp) into the pPD95.83 vector. Worms were raised with the HT115 strain of Escherichia coli, and RNAi was performed as described. TMRE experiments were performed by synchronizing and raising worms on plates previously soaked with M9 buffer containing 10μM TMRE (Mallick and Haynes, 2024).

### Analysis of worm development

Worms were synchronized via bleaching and allowed to develop on HT115 bacteria plates at 20°C. The developmental stage was quantified as a percentage of the total number of animals that turned adult after an incubation of 58 hrs at 20°C. Each experiment was performed three times. For the comparison of wild-type, *atfs-1(null)* and *atfs-1(null); nhr-180*(RNAi) worms; N = 162 (wild type), and 282 (*atfs-1(null)*). For the comparison of wild type to *isp-1*(mutant) and *clk-1*(mutant) worms, N = 158 (wild type) and N = 256 (*isp-1*).

### Starvation assay

Worms were bleach-synchronized, and eggs were plated on HT115 *E. coli*. Worms grew at 25°C until bacterial depletion induced dauer formation. Dauer larvae were then transferred to fresh bacterial plates to exit arrest. This cycle was repeated two additional times before worms were bleached again and analyzed for lifespan and TMRE staining at the L4 stage.

### Pharyngeal pumping and locomotion

The pharyngeal pumping assay was performed on NGM plates containing a thin bacterial lawn. Worms were bleach synchronized and allowed to grow in the presence of food to the late L4 stage. 10 worms were transferred to assay plates, and the number of contractions in the terminal bulb of the pharynx of each animal was counted every 24 hours for 30 sec using a Nikon 80i inverted microscope. Four independent experiments were performed each day. The locomotion rate of worms on different days of adulthood was examined using a protocol from the Sternberg lab. Briefly, worms were bleach-synchronized and allowed to grow till the late L4 stage. Five worms were placed onto separate plates and tested daily for locomotion until death. For testing, a worm was placed onto an NGM plate containing a uniform layer of bacteria and stimulated by contact with the tail. The number of body movements was counted from the trail left on the plate (Mallick et al., 2020)

### Lifespan assay

Lifespan analysis was carried out following an established protocol (Amrit et al., 2014; Mallick et al., 2020). Each strain was repeated at least twice. At least 50 animals were used per condition, and worms were scored for viability every second day, from day 1 of adulthood (treating the pre-fertile day preceding adulthood as t = 0). Young adult worms were transferred to fresh plates every other day, and the numbers of dead worms were recorded as events scored. Animals that were lost or burrowed in the medium, exhibiting protruding vulva (intestine protrudes from the vulva), or undergoing bagging (larvae hatching inside the worm body) were censored. Prism 8 (GraphPad) software was used for statistical analysis, and *p-values* were calculated using the log-rank (Kaplan-Meier) method.

### Protein analysis and antibodies

Synchronized worms were raised on plates with control(RNAi) or *cco-1*(RNAi) to the L4 stage before harvesting. The whole worm lysate preparation was previously described33. Antibodies against β-actin (cell signaling), anti-FLAG M2 antibody (Sigma, F1804), MTCO1 (CTC-1 in *C. elegans*, Abcam), and NDUFS3 (NUO-2 in *C. elegans*, complex I, Abcam Invitrogen) were used. Polyclonal antibodies were generated against amino acids 953–971 of *C. elegans* LONP-1 and subsequently affinity-purified by Thermo Fisher Scientific. All antibodies were diluted 1:2000. Immunoblots were visualized using the ChemiDoc XRS + system (Bio-Rad). All western blot experiments were performed at least three times.

### Cellular thermal shift assay (CETSA)

For each assay, ∼100,000–200,000 L4 3xFLAG::NHR-180, 3xFLAG::NHR-86, or NHR-12::3xFLAG worms were lysed in PBS with HALT protease inhibitors using a Teflon Dounce homogenizer. Lysates were clarified by centrifugation (16,000 rpm, 20 min, 4°C), quantified via DC Protein Assay (Bio-Rad), and adjusted to 10 mg/mL. Aliquots were treated with water or 100 µM of AMP or ATP for 15–60 minutes at room temperature. During incubation, 50 µL of each lysate was exposed to 25–65°C for 4 minutes, cooled, and chilled on ice. After centrifugation (20,000 g, 20 min, 4°C), supernatants were mixed with LDS Sample Buffer and 1% β-mercaptoethanol (Peterson et al., 2023; Tse and Pukkila-Worley, 2023).

### mtDNA quantification

mtDNA quantification was performed using a quantitative PCR (qPCR)-based method similar to previously described assays (Lin et al., 2016; Yang et al., 2022). Twenty to 30 worms were collected in 30 μl of lysis buffer (50mM KCl, 10mM Tris-HCl pH 8.3, 2.5mM MgCl2, 0.45% NP-40, 0.45% Tween 20, 0.01% gelatin, with freshly added 200 μg/ml proteinase K) and frozen at −80 °C for 20 min before lysis at 65 °C for 80 min. Relative quantification was used to determine the fold changes in mtDNA between samples. 1 μl of lysate was used in each triplicate qPCR reaction. qPCR was performed using the Thermo Scientific SyBr Green Maxima Mix and the MyiQ2 Two-Color Real-Time PCR Detection System (Bio-Rad Laboratories). Primers that amplify a non-coding region near the nuclear-encoded ges-1 gene were used as a control. mtDNA was harvested from synchronized worms at the L4 stage. All qPCR results have been repeated at least three times and performed in triplicate. A Student’s t-test was employed to determine the level of statistical significance.

### Cell culture

The KSS cell line was a gift from C. Moraes (Moraes et al., 1989). SDHA KO cells were obtained from Jessica Spinelli’s lab (Spinelli et al., 2021). ATF5(-) cell line was created in the lab using CRISPR-Cas9. Cells were cultured in DMEM (4 mM l-glutamine, 4.5 g per liter glucose; Gibco, Thermo Fisher Scientific) plus 10% FBS with 1% penicillin-streptomycin. Total cellular mtDNA was prepared as previously described (Fiorese et al., 2016). Cells were incubated continuously in the described concentration of CDDO for the indicated number of days. The cells were subcultured before confluence every 48 h.

### RNA isolation and qRT-PCR

RNA isolation and quantitative reverse transcriptase PCR (qRT-PCR) analysis were previously described (Lin et al., 2016). Worms were synchronized by bleaching, raised on HT115 *E. coli,* and harvested at the L4 stage. Total RNA was extracted from frozen worm pellets using Trizol reagent, and 500 ng RNA was used for cDNA synthesis with qScript™ cDNA SuperMix (QuantaBio). qPCR was performed using iQ™ SYBR® Green Supermix (Bio-Rad Laboratories). All qPCR results were repeated at least 3 times and performed in triplicate. A two-tailed Student’s t-test was employed to determine the level of statistical significance.

### Microscopy

*C. elegans* were imaged using either a Zeiss AxioCam 506 mono camera mounted on a Zeiss Axio Imager Z2 microscope or a Zeiss AxioCam MRc camera mounted on a Zeiss SteREO Discovery.V12 stereoscope. Images with high magnification (×63) were obtained using the Zeiss Apotome 2. Exposure times were the same in each experiment. Cell cultures were imaged with the Zeiss LSM800 microscope. With Zen 2.3 Blue software. All images are representative of more than three images. Quantification of fluorescent intensity as well as creating binary skeleton-like structures was done with ImageJ.

### siRNA

Cells were grown in 6-well plates, and siRNAs were transfected with Lipofectamine RNAiMAX (Thermo Fisher Scientific, 13778150) following the manufacturer’s instructions. Human *S6K* siRNA was a gift from Dr. Claudio Punzo’s lab.

### Respiration assays

#### Cell line

For mitochondrial respiration assays, the OCR was measured using a Seahorse Extracellular Flux Analyzer XFe96 (Seahorse Biosciences) as previously described (Fiorese et al., 2016). A total of 14,000 CoxI G6930A cells were seeded per well with fresh medium treated with DMSO or a defined concentration of S6K inhibitor PF-4708671 (Selleckchem, USA). The OCR was measured using a Cell MitoStress kit (as described by the manufacturer). A volume of 180 μl of XF-Media was added to each well, and then the plates were subjected to analysis following sequential introduction of 1.5 μM oligomycin, 1.0 μM FCCP, and 0.5 μM rotenone/antimycin as indicated. Data is normalized to total protein as determined by the BCA protein assay.

#### Worms

Oxygen consumption was measured using a Seahorse XFe96 Analyzer at 25°C, similar to that described previously (Koopman et al., 2016). In brief, L4 worms were transferred onto empty plates and allowed to completely digest the remaining bacteria for 1 h, after which ten worms were transferred into each well of a 96-well microplate containing 180 μl M9 buffer. Basal respiration was measured for a total of 30 min, in 6 min intervals that included a 2 min mix, a 2 min time delay, and a 2 min measurement. To measure respiratory capacity, 15 μM carbonyl cyanide-4-(trifluoromethoxy)phenylhydrazone was injected, the oxygen consumption rate reading was allowed to stabilize for 6 min and then measured for five consecutive intervals. Mitochondrial respiration was blocked by adding 40 mM Sodium azide. Each measurement was considered one technical replicate.

### Molecular modeling and molecular dynamics simulations

AlphaFold 3 was used to predict the structure of the full-length NHR-180 (Abramson et al., 2024). To determine an optimal binding pocket, AutoDock Vina and ClusPro software were used (Eberhardt et al., 2021; Kozakov et al., 2017). The potential interaction between each ligand of interest and NHR-180 was done using AutoDock Vina and ClusPro and hydrogen bonding network was determined. Structural figures were generated using PyMOL (v. 2.3.4).

### RNA-seq and differential expression analysis

cDNA libraries were constructed with standard Illumina P5 and P7 adapter sequences. The cDNA libraries were run on an Illumina HiSeq2000 instrument with single-read 50 bp. RNA reads were then aligned to the WS295 reference genome, and differential gene expression analysis was performed with edgeR43. Differences in gene expression between *nhr-180(cmh19), atfs-1(null),* and *atfs-1(null); nhr-180(cmh19)* compared to wild type are listed in Supplementary Table 2, respectively. Data were deposited in the GEO accession no. GSE290561.

### Gene set enrichment analysis

The gene set was downloaded from the WormBase Ontology Browser. mRNA abundance was measured and ranked by reads per kilobase per million reads from RNA-seq data. Pre-ranked gene set enrichment analysis was performed with GSEA3.0 software with ‘classical’ scoring.

### Statistics

All experiments were performed at least three times, yielding similar results and comprised of biological replicates. The sample size and statistical tests were chosen based on previous studies with similar methodologies, and the data met the assumptions for each statistical test performed. No statistical method was used in deciding sample sizes. No blinded experiments were performed, and randomization was not used. For all figures, the mean ± SD is represented unless otherwise noted. Prism 8 (GraphPad) is used for statistical analysis and graph creation.

## Data availability

The data reported in this paper have been deposited in the Gene Expression Omnibus (GEO) database (accession number GSE290561). They are also available from the corresponding author upon reasonable request. Source data is provided in this paper.

## Acknowledgments

We thank the *Caenorhabditis* Genetics Center for providing *C. elegans* strains (funded by NIH Office of Research 362 Infrastructure Programs (P40OD010440), and the UMass Medical School Core facilities for deep sequencing. This work was supported by the National Institutes of Health grants (R01AG040061 and R01AG047182) to C.M.H. and the Natural Sciences and Engineering Research Council (NSERC) postdoctoral fellowship to A.M. The authors are solely responsible for the content. We are grateful to the Pukkila-Worley lab for all the support throughout the project.

## Author contributions

A.M. and C.M.H. planned the experiments. A.M. and Y.D. generated worm strains. A.M. performed RNA-seq, lifespan analysis, qPCR, and TMRE experiments. T.R. carried out experiments in the mammalian cell lines. R.L., and L.J.Z. analyzed the sequencing data. L.A. and S.K. performed some of the experiments. T.R. performed mammalian cell culture assays. P.R. performed the ligand binding prediction. A.M. and C.M.H. wrote the manuscript.

## Supplementary files

### Supplementary Figures

**Supplementary Figure 1:**
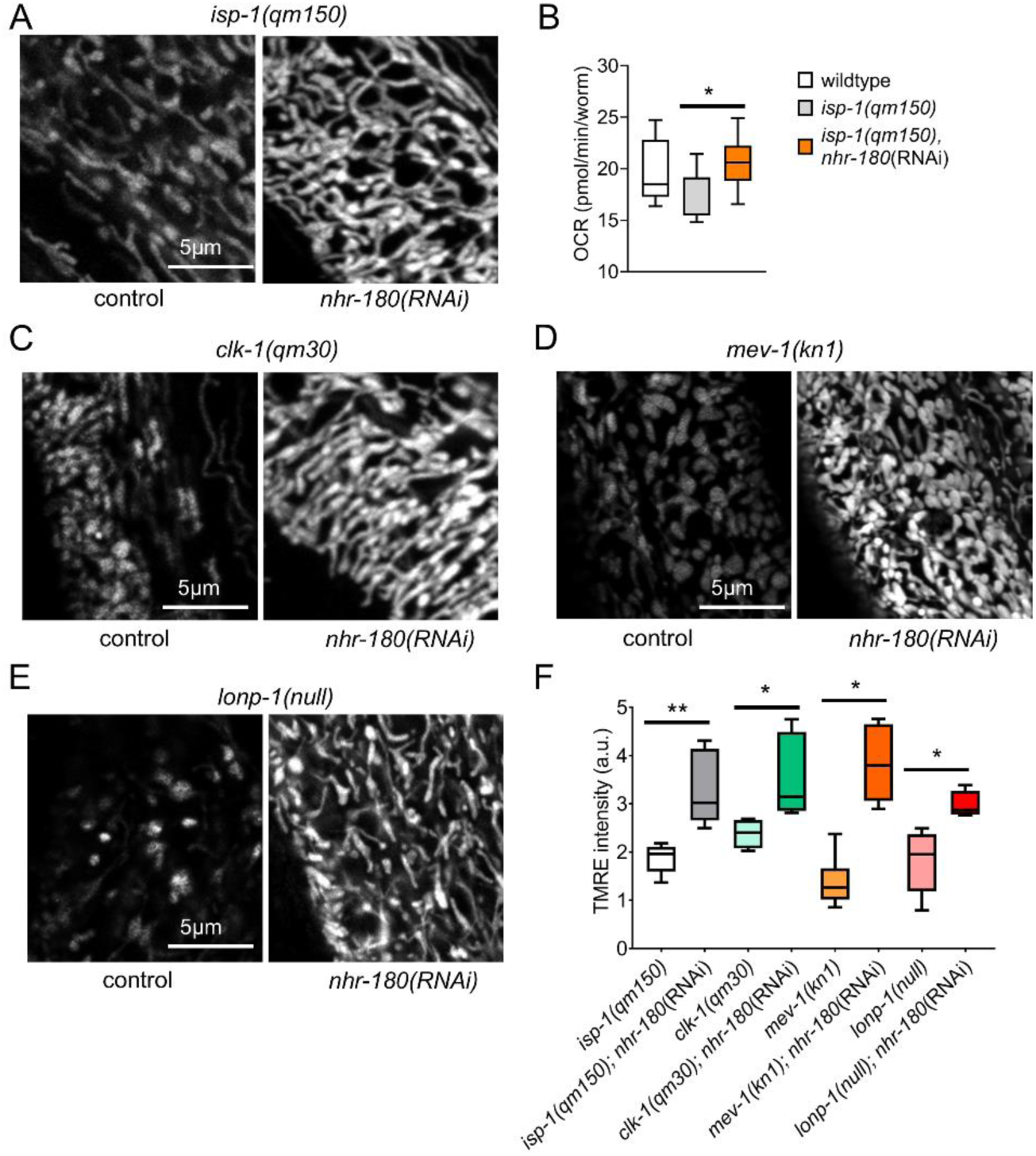
Inhibition of *nhr-180* increases mitochondrial function in the OXPHOS mutants. **A** TMRE staining of *isp-1(qm150)* worms after control or *nhr-180*(RNAi). The scale bar represents 5μm. Experiments were repeated three times with similar results. **B** Oxygen consumption rate (OCR - pmol/min) in wildtype, *isp-1(qm150)* and *isp-1(qm150); nhr-180*(RNAi) worms at the L4 stage. Data is plotted as the average of the three replicates, and error bars indicate SEM. * *p* < 0.05 using two-tailed unpaired t-test. **C-E** TMRE staining of *clk-1(qm30), mev-1(kn1)* and *lonp-1(null)* worms after control or *nhr-180*(RNAi). The scale bar represents 5μm. Experiments were repeated three times with similar results. **F** Quantification of TMRE intensity for experiments in **A, C-E**. Data is plotted as the average of the three replicates, and error bars indicate SEM. * *p* < 0.05 and ** *p* < 0.01 using two-tailed unpaired t-test.

**Supplementary Figure 2:**
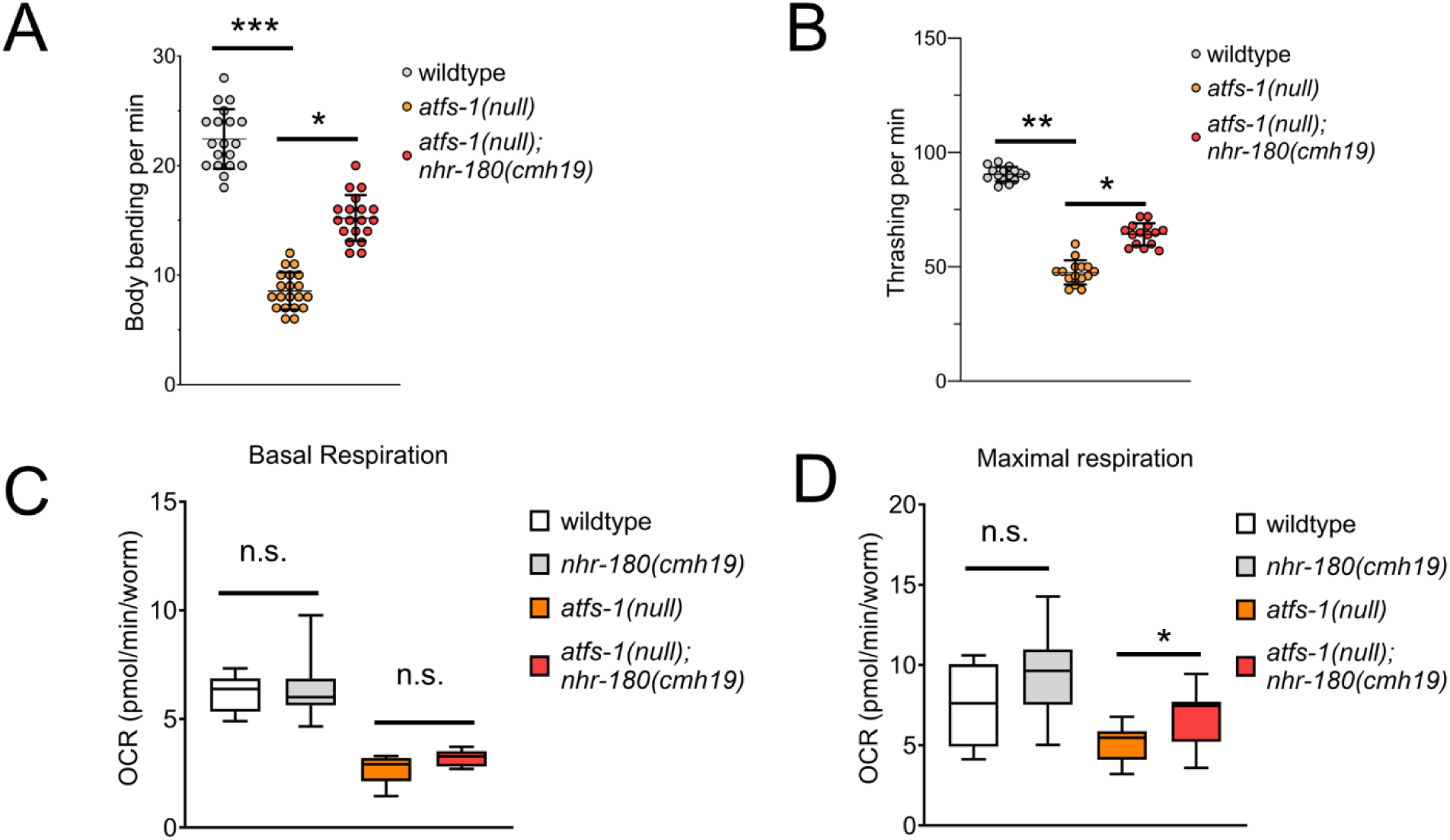
Mutation in *nhr-180* leads to increased body bending and thrashing rate of *atfs-1(null)* worms. **A,B** Dot plots showing the body bending and thrashing rates of wildtype, *atfs-1(null)*, and *atfs-1(null); nhr-180(cmh19)* adult worms. Biologically, 3 independent samples, * *p* < 0.05, ** *p* < 0.01, and *** *p* < 0.0001 (multiple unpaired t-test). **C, D** Box, and violin plots showing the basal and maximal respiration of wildtype, *nhr-180(cmh19), atfs-1(null)*, and *atfs-1(null); nhr-180(cmh19)* worms at the L4 stage. Biologically, 2 independent samples, ** *p* < 0.01 (two-tailed unpaired t-test). Graphs represent the average of the replicates, with error bars indicating the SEM.

**Supplementary Figure 3:**
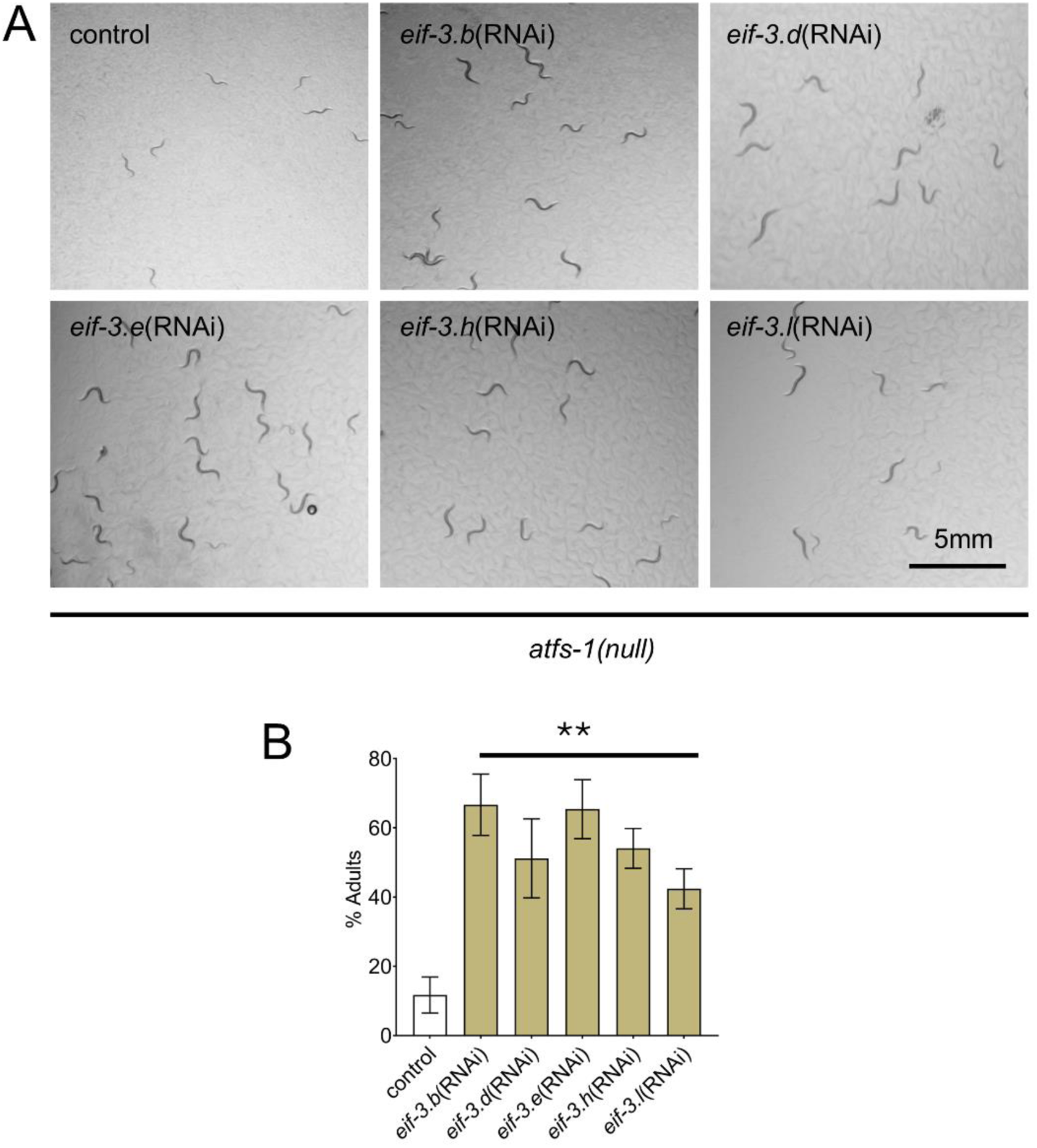
*atfs-1(null)* worms develop faster following inhibition of EIF-3 complex components. **A** Representative image showing *atfs-1(null)* worms at the plate level following inhibition of *eif-3.b*, *eif-3.d*, *eif-3.e*, *eif-3.h* and *eif-3.l* for 58 hrs. The scale bar represents 5mm. **B** Bar graph showing the percentage of *atfs-1(null)* worms that turned into adults after an incubation for 58 hrs at 20°C following RNAi inhibition shown in **A**. Biologically, 3 independent samples, ** *p* < 0.01 (multiple unpaired t-test). Bar graphs represent the average of the replicates, with error bars indicating the SEM.

**Supplementary Figure 4:**
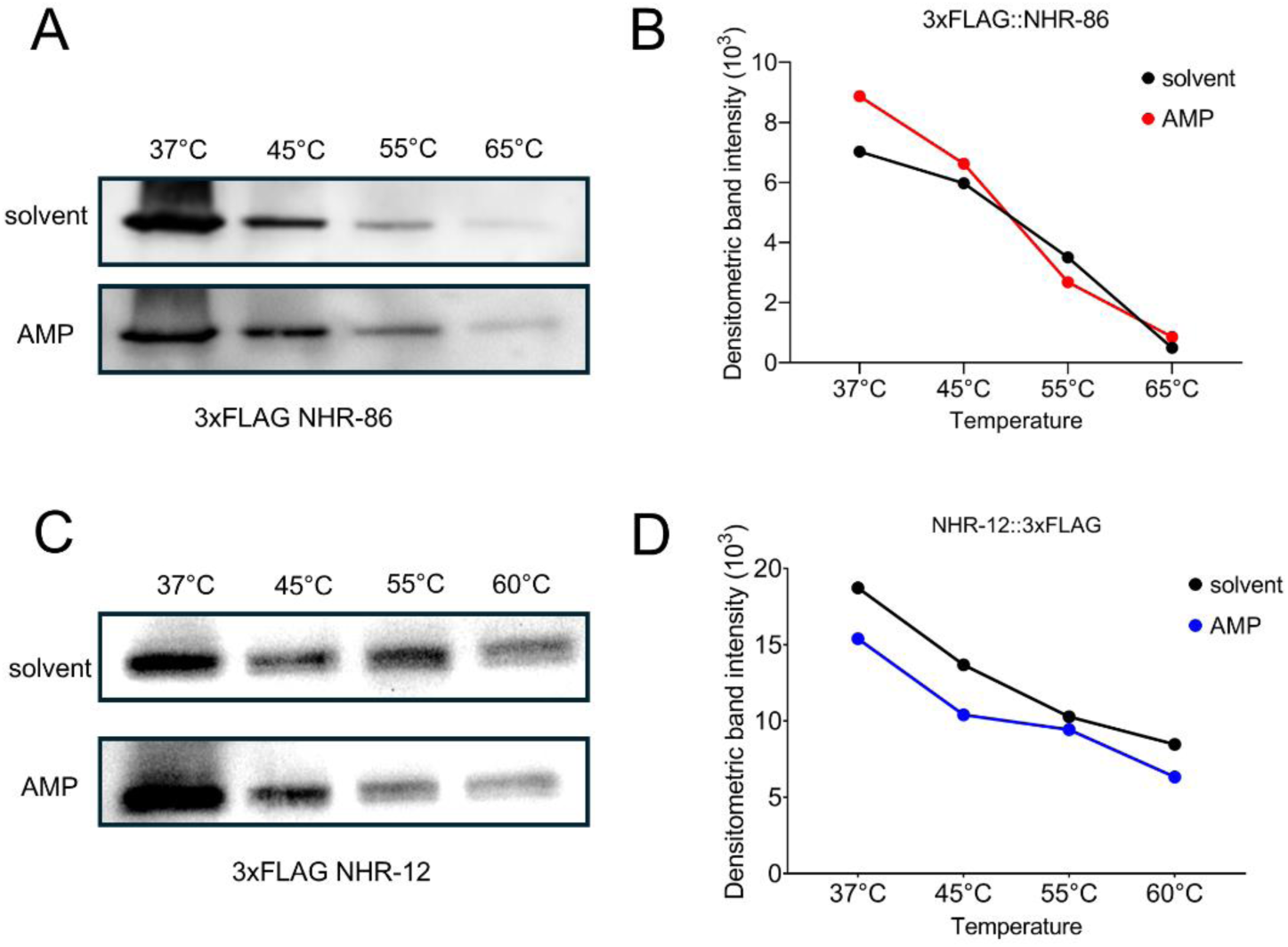
AMP does not stabilize NHR-86 or NHR-12 proteins. **A,C** A representative immunoblot of a cellular thermal shift assay (CETSA) experiment using an anti-FLAG antibody that probed whole-cell lysates from a transgenic *C. elegans* strain in which NHR-86 or NHR-12 was tagged with 3xFLAG at its endogenous locus. **B, D** A representative densitometric quantification from a CETSA experiment that characterized the interaction of AMP (100µM), and solvent (water) with 3xFLAG::NHR-86 or NHR-12::3xFLAG.

**Supplementary Figure 5:**
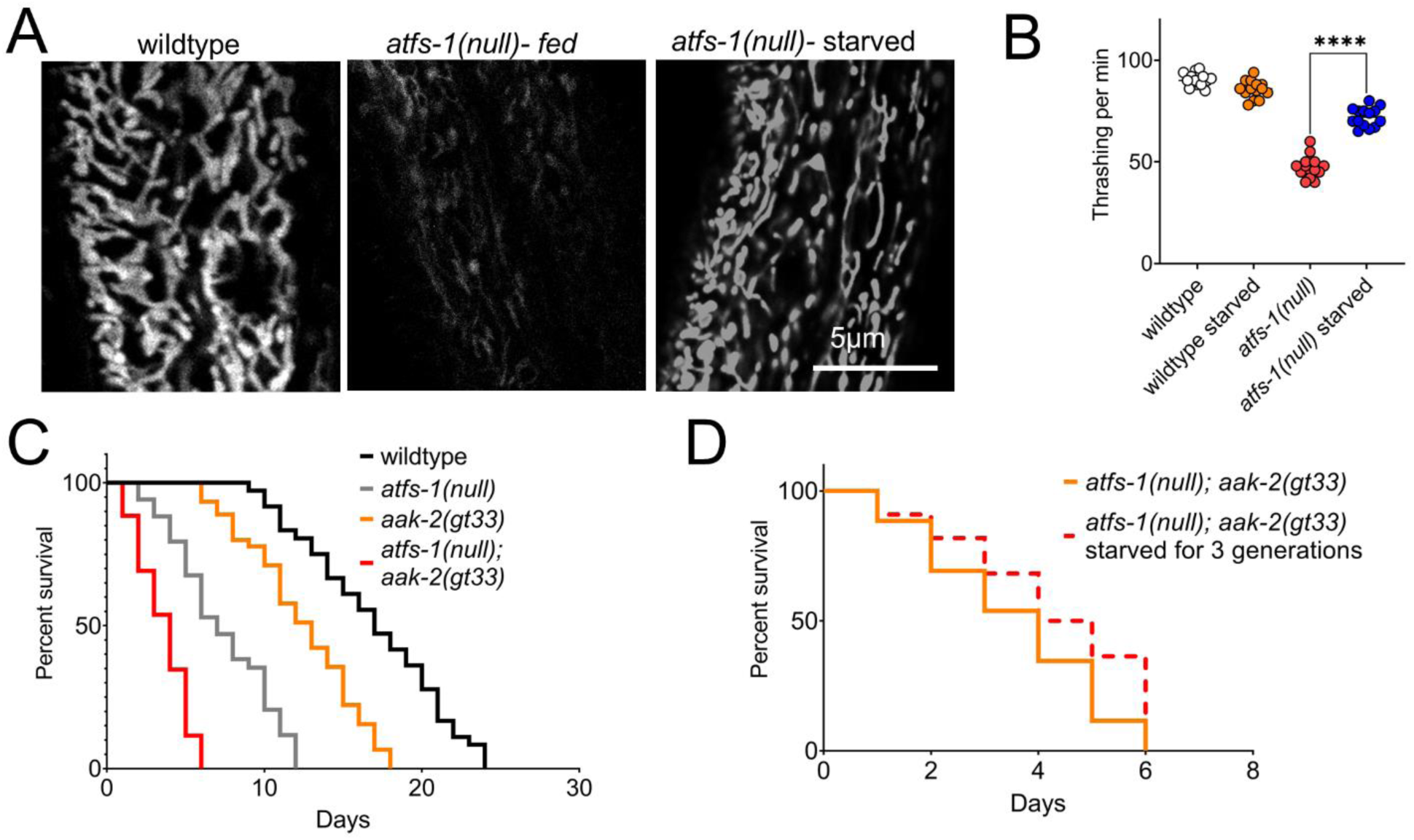
Starvation increases mitochondrial function. **A** Representative image of TMRE staining on L4 worms, scale bar represents 5µm. **B** Dot plots represent the thrashing rate per minute for wildtype and *atfs-1(null)* day 1 adults after feeding and starvation (See Methods for the starvation protocol). **C** Lifespan analysis for wildtype, *atfs-1(null)*, *aak-2(gt33)* and *atfs-1(null); aak-2(gt33)* worms. **D** Lifespan analysis of *atfs-1(null); aak-2(gt33)* double mutant worms in the presence and absence of starvation. **C, D** N =3, biologically independent replicates. **(C)** \**p* < 0.01 and **(D)** not significant (log-rank test).

### Supplementary Tables

**Supplementary Table 1:** List of transcription factors RNAi in *atfs-1(null)*.

**Supplementary Table 2:** List of differentially expressed genes in *nhr-180(cmh19)* vs wildtype.

**Supplementary Table 3:** List of cytosolic protein translation genes downregulated in *nhr-180(cmh19)* worms.

**Supplementary Table 4:** List of mitochondrial protein translation genes downregulated in *nhr-180(cmh19)* worms.

**Supplementary Table 5:** List of larval development genes that are upregulated in the *nhr-180(cmh19)* worms.

**Supplementary Table 6:** List of common upregulated genes between *nhr-180(cmh19)* and *eat-2(ad465)* worms.

**Supplementary Table 7:** List of common upregulated genes between *nhr-180(cmh19)* and AAK-2(OE) worms.

